# No trace of phase: Corticomotor excitability is not tuned by phase of pericentral mu-rhythm

**DOI:** 10.1101/513390

**Authors:** Kristoffer Hougaard Madsen, Anke Ninija Karabanov, Lærke Gebser Krohne, Mads Gylling Safeldt, Leo Tomasevic, Hartwig Roman Siebner

## Abstract

**Background:** The motor potentials evoked by transcranial magnetic stimulation (TMS) over the motor hand area (M1-HAND) show substantial inter-trial variability. Pericentral mu-rhythm oscillations, might contribute to inter-trial variability. Recent studies targeting mu-activity based on real-time electroencephalography (EEG) reported an influence of mu-power and mu-phase on the amplitude of motor evoked potentials (MEPs) in a preselected group with strong pericentral mu-activity. Other studies that determined mu-power or mu-phase based on post-hoc trial sorting according in non-preselected individuals were largely negative.

**Objectives:** To reassess if cortico-spinal activity is modulated by the mu-rhythm, we applied single-pulse TMS to the M1-HAND conditional on the phase of the intrinsically expressed pericentral mu-rhythm in 14 non-preselected healthy young participants.

**Methods:** TMS was given at 0, 90, 180, and 270 degrees of the mu-phase. Based on the absence of effects of mu-phase or mu-power when analyzing the mean MEP amplitudes, we also computed a linear mixed effects model, which included mu-phase, mu-power, inter-stimulus interval (ISIs) as fixed effects, treating the subject factor as a random effect.

**Results:** Mixed model analysis revealed a significant effect of mu-power and ISI, but no effect of mu-phase and no interactions. MEP amplitude scaled linearly with lower mu-power or longer ISIs, but these modulatory effects were very small relative to inter-trial MEP variability.

**Conclusion:** Our largely negative results are in agreement with previous offline TMS-EEG studies and point to a possible influence of ISI. Future research needs to clarify under which circumstances the responsiveness of human the M1-HAND to TMS depends on the synchronicity with mu-power and mu-phase.

**Highlights:**

- Phase-triggered TMS at four distinct phases of the ongoing mu-oscillations is technically feasible in non-preselected young volunteers
- Targeting the ongoing mu-activity did not reveal consistent modulatory effect of mu-phase on corticospinal excitability in a non-preselected group
- Mixed-effects analysis revealed a weak but significant effect of pre-stimulus mu-power and ISI on corticospinal excitability

## Introduction

Cortical oscillations play an important role in information processing in brain networks [1-3]. The occipital alpha rhythm (8-12 Hz) is a prominent oscillatory signature, and regional variations in posterior alpha power have been proposed to gate visual processing by active inhibition of task-unrelated areas [4-7]. The pulsed-inhibition hypothesis postulates that the occipital alpha rhythm creates periods of inhibition depending on the oscillation phase. The peak of the alpha oscillation is characterized by a high level of inhibition, while there is a low level of inhibition during the trough of the alpha oscillation, providing a preferred window for visual information processing [8, 9]. Studies using single-pulse Transcranial Magnetic Stimulation (TMS) have also provided evidence in favor of inhibitory modulation and have shown that a single TMS pulse over the occipital cortex has a stronger likelihood of eliciting an illusionary percept (i.e. phosphenes) when given during periods of low alpha power [10, 11]. Accordingly, psychophysiological experiments showed power- and phase-dependent modulation of visual perception [12, 13].

The pericentral Rolandic cortex also expresses prominent oscillatory activity in the alpha range, commonly referred to as the mu-rhythm [14]. The pericentral mu-rhythm has been shown to modulate the perception of somatosensory stimuli in a manner similar to the modulation of visual perception by occipital alpha [15-18]. Cortical *in vivo* recordings in monkeys revealed that pericentral alpha activity modulates the normalized firing rate in the sensorimotor regions. In agreement with the pulsed inhibition hypothesis, these data revealed higher firing rates at the trough and lower firing rates at the peak of the alpha oscillations. Cortical firing rates were also modulated by fluctuations in alpha power. Firing rate was reduced when alpha activity was weak (low alpha power) relative to epochs with prominent alpha activity (high alpha power), indicating an inverse relationship between alpha power and cortical neural activity [19].

Single-pulse TMS of the motor hand area (M1-HAND) has been combined with electroencephalography (EEG) to test how ongoing pericentral oscillatory activity impacts corticomotor excitability as reflected by the amplitude of the motor evoked potential (MEP) [20, 21]. Most studies have adopted a “offline approach”, using posthoc trial sorting to test whether EEG power or phase of the Rolandic mu-rhythm scales with the MEP amplitude [22-31]. The results of these studies were largely negative and are summarized in table 1. Two early studies found a negative linear relationship between pre-stimulus mu-power and the MEP amplitude at rest [22, 23] as predicted by the invasive recordings in monkeys and the pulsed inhibition hypothesis [19]. The same relationship was also reported by a study in the context of event-related desynchronization [26]. All later post-hoc sorting studies consistently failed to replicate a modulatory impact of pre-stimulus mu-power on MEP amplitude (Table 1). A single study reported a relationship between pre-stimulus variability in alpha power (but not magnitude of power) and variability in MEP amplitude [30]. Some studies rather observed associations of MEP amplitude with the pericentral cortical or cortico-muscular expression of beta band activity (Table 1).

**Table 1:**
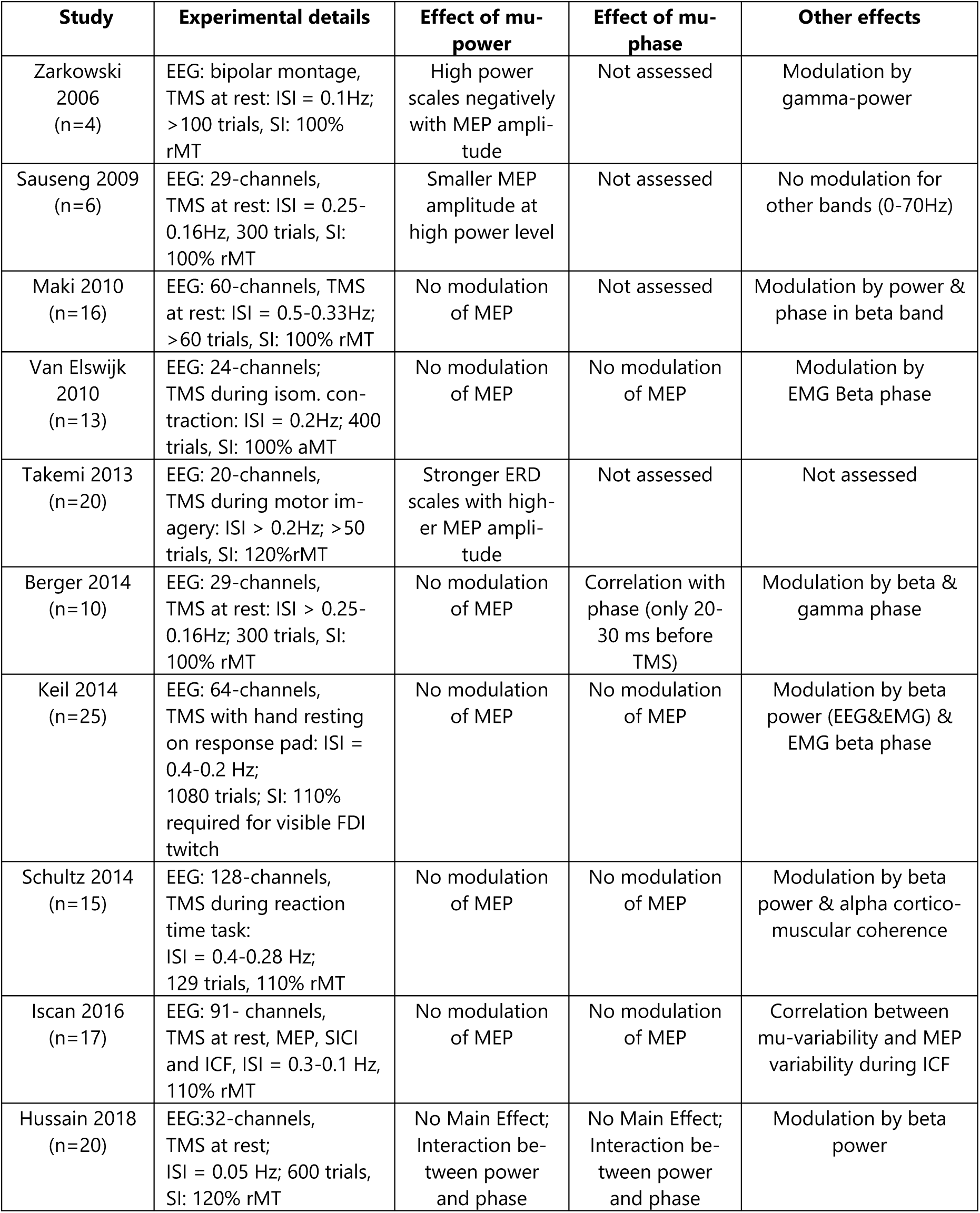
Synopsis of offline TMS-EEG studies that have used post-hoc trial sorting to investigate relations between mu-phase and/or power and MEP amplitude. All studies have used focal TMS to target the M1-HAND. ISI = Inter-stimulus interval (ISI), SI = stimulation intensity of TMS, rMT = resting motor threshold, aMT = active motor threshold.

Evidence for a modulatory impact of phase on the MEP amplitude is even scarcer; only one study reported a relationship between the alpha oscillation phase 30-40ms before the TMS pulse and the MEP size [27], others did not find mu-phase-dependent MEP modulations [25, 27-29, 31]. Using a mixed-model analysis, a recent study reported a flip in the relationship between mu-phase and MEP amplitude depending on the level of mu-power at the time of TMS [31]: When alpha power was low, MEP amplitudes were lower during the trough of the mu-oscillation, whereas MEP amplitudes were higher during the trough of the mu-rhythm when power was high. No clear relationship between mu-phase and MEP amplitude was present at medium levels of alpha power [31].

In recent years, an online strategy has been successfully established which delivers the TMS pulse based on the real-time EEG expression of the target oscillation. EEG-informed phase-dependent TMS was first applied during non-REM sleep [32] and recently applied during resting wakefulness. In highly pre-selected groups of healthy individuals with strong pericentral mu-activity, single-pulse TMS was applied to M1-HAND dependent on mu-phase [33-35] or mu-power [36] of the locally expressed mu-activity. These studies found higher MEP amplitudes at the trough relative to the peak of the mu-phase [33, 34] as well as higher MEP amplitude in epochs with high relative to low mu-power [36], while no interaction between the two was reported. While the phase dependent modulation is in line with the invasive recordings in monkeys [19], the power modulation is opposite in sign, suggesting a “pulsed facilitation” rather than a “pulsed inhibition” mechanism.

In this study, we re-assessed the influence of mu-phase and mu-power on corticospinal excitability. We applied a brain-state informed EEG-TMS method for real-time phase estimation of the mu-rhythm that does not require any pre-selection of subjects based on the magnitude of their endogenous mu activity [37]. We targeted not only the peak (0°) and trough (180°), but also the time points of steepest increase (90°) and steepest decrease (270°) of the mu-oscillations. In addition to mu-phase, we tested whether mu-power and the time that elapsed between two consecutive TMS pulses influenced MEP amplitude and its modulation by mu-phase.

## Methods

### Subjects

14 healthy volunteers were recruited to take part in the study (5 female, average age: 22.9 y ± 2.3). Participants were *not* pre-selected based on individual TMS or EEG characteristics such as motor resting threshold (RMT) or the presence of a clear alpha peak in the power spectrum over the sensorimotor cortex at rest. All subjects gave written consent and the study was approved by the Regional Committee on Health Research Ethics of the Capitol Region in Denmark in accordance with the declaration of Helsinki (Protocol H-16017716). Sample size was based on previous studies investigating instantaneous mu-phase modulations of corticospinal excitability [34].

### Experimental set-up

Throughout the experiment, participants were sitting in a relaxed position in a commercially available TMS-chair (MagVenture, Farum, Denmark). Additional cushioning provided additional arm and neck support and the participant was instructed to keep the hands and arms relaxed and the eyes open throughout the experiment.

We used a real-time EEG-TMS setup for online analysis of the endogenously expressed EEG-signal and triggered single TMS pulses at a specific phase of the intrinsically expressed pericentral mu-rhythm (see Figure 1). Single-pulse TMS was performed with a figure-of-eight shaped MC-B70 coil connected to a MagPro 100 stimulator (Magventure, Farum, Denmark). TMS targeted the left M1-HAND using monophasic pulses triggered by an external trigger pulse generated by the real-time signal processing system. Stimulation intensity was individually adjusted to elicit a mean MEP amplitude of approximately 1 mV (see experimental procedure for details). MEPs were recorded from the completely relaxed first dorsal interossus (FDI) muscle of the right hand using self-adhesive, disposable surface electrodes (Neuroline, Ambu A/S, Denmark) in a belly-tendon montage. The motor hotspot was defined as the coil location and orientation that elicited the largest MEP amplitude in the relaxed FDI representation. The FDI motor hotspot was the individual target site and precise positioning of the TMS coil on the target was continuously monitored with the help of stereotactic neuronavigation (Localite GmbH, Sankt Augustin, Germany). Neuronavigation was also used for recording the position of the EEG electrodes relative to the individual brain as mapped with structural MRI.

**Figure 1:**
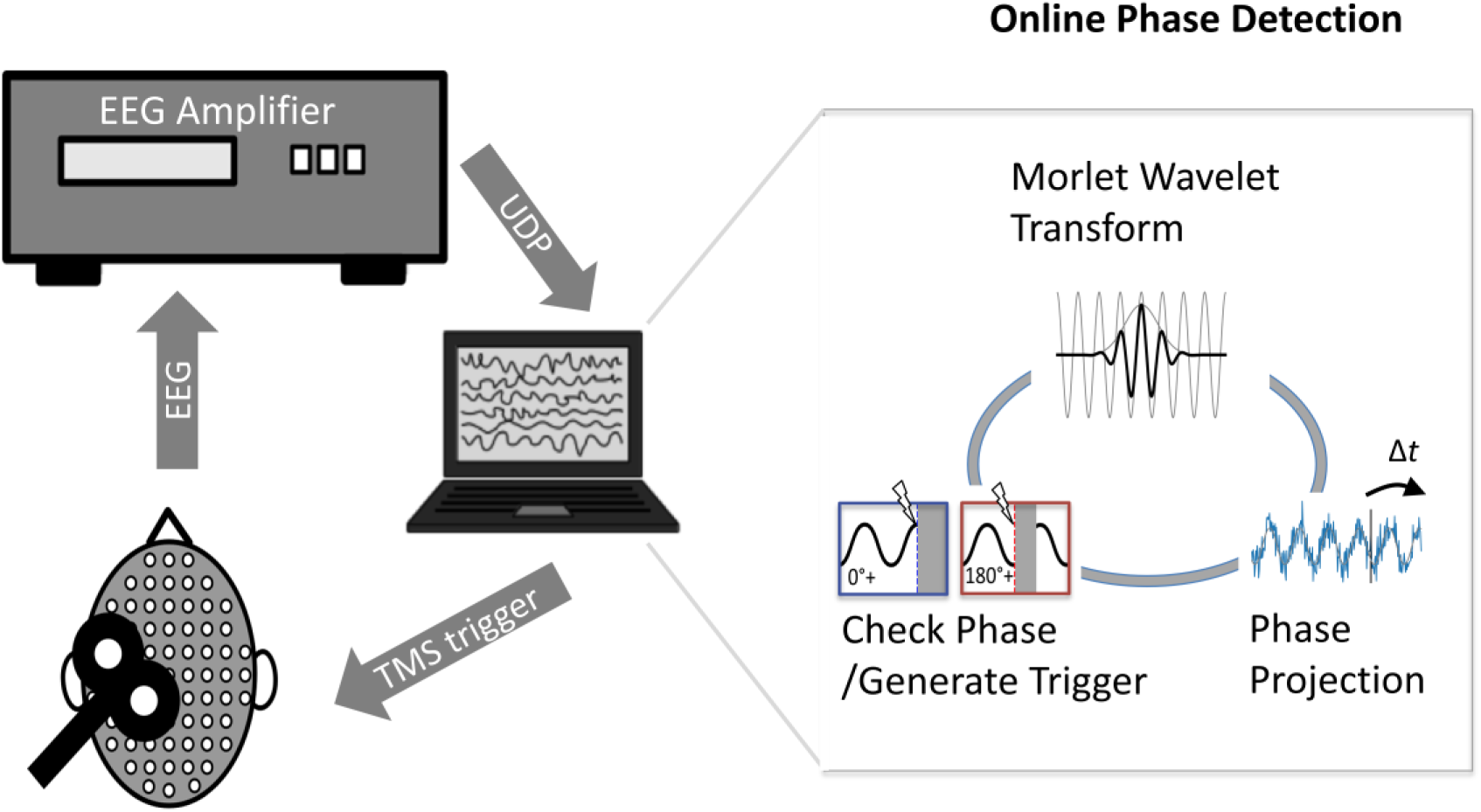
Brain state informed EEG-TMS setup for online phase detection and phase targeting with TMS. Based on the extracted EEG signal, the phase is forward projected to the current time. TMS will be triggered, if the current estimated phase is close to the target phase.

Resting motor threshold (MEP ≥ 50uV) and test intensity (MEP ≥ 1mV) were determined using an adaptive threshold-hunting algorithm (Groppa S *et al*. 2012). Threshold hunting was initiated at 47% of maximum stimulator output (MSO) and the relative standard deviation of the true threshold was assumed to be 7% during threshold hunting.

### Electrophysiological recordings

Scalp EEG was recorded from a 63-channel TMS compatible, equidistant ring electrode cap (Easycap M10 sintered Ag/AgCl multitrodes EasyCap, Woerthsee-Etterschlag, Germany). EEG and MEP were recorded using a Bittium NeurOne Tesla EEG amplifier utilizing 5 kHz sampling rate and a 2.5 kHz antialiasing lowpass filter with 24-bits resolution per channel across a range of ±430 mV (NeurOne Tesla, Bittium, Oulu, Finland). For real-time processing, data was sent directly from the EEG amplifier Field-Programmable-Gate-Array (FPGA) via user-datagram-protocol (UDP) packets, each consisting of five samples, resulting in a 1 kHz update rate over a 1 Gb/s Ethernet link. This setup ensured that the total latency of packages delivered to the real-time analysis loop could be kept below 5 ms.

### Real Time Digital Signal Processing

During the experiment, the instantaneous phase of the mu-rhythm was continuously estimated to enable phase targeting using in-house developed analysis software. This software was implemented in Python 2.7 and consisted of two parts, each running as independent processes on a standard computer. 1) A data receive process collecting the data from the amplifier via UDP, and continuously updating a ring-buffer with the last 500 ms of data. 2) A phase estimation and stimulation loop. The division into separate processes ensured that no packets were dropped in the data collection and that phase estimated could be performed with minimal latency. The phase estimation process was running asynchronously at a best effort update rate, where each loop started by requesting the latest data window for all channels from the data receive process. The data from the 63-EEG channels was then projected to source space (see Section on Experimental Session for details), linearly detrended, and the phase was estimated using a continuous wavelet transform within the 500 ms window.

Continuous phase estimates are in principle possible for all time points within the time windows, but the estimates become severely distorted near the edges of the window. Therefore, we used the estimated phase and frequency at 140 ms prior to the end of the window to project the phase to the current time point. We choose that time point based on simulated data, indicating that this position within the time window provided a reasonable trade-off between immediate online targeting and estimation accuracy [37]. The wavelet transform was based on the continuous Morlet mother wavelet within 51 frequency scales uniformly distributed across the desired frequency range. To improve the computational efficiency of the estimation, the Fourier transformed wavelet basis functions were pre-calculated and the inverse fast Fourier transform using the FFTW library (version 3.3.5) [38] considered only frequencies up to 500 Hz. A stimulation trigger was generated and sent via a standard parallel port interface if the following three criteria were met. 1) Estimated phase was within ±10° of the intended stimulation phase. 2) The power within the considered mu frequency band was above the 75th percentile, as estimated from a separate rest EEG recording immediately prior to the main experiment. 3) The phases within the latest 100 ms did not differ from the previous estimates by more than 90°. The last criterion served to ensure that stimulation did not occur when the phase estimation was unstable, for instance in the presence of artefacts caused by eye blinks or other muscle movement. Due to the efforts to improve the computational efficiency of the phase estimation loop the update loop was able to run at an average update frequency of approximately 1kHz, adding only a negligible amount of additional latency to the setup.

### Experimental sessions

#### Structural scans

To be able to record the individual position of EEG electrodes and to monitor coil placement throughout the experiment each participant underwent a structural T1-weighted MRI scan using a magnetization prepared rapid gradient echo (MPRAGE) sequence with TR = 6 ms, TE = 2.70 ms; flip-angle = 8°, 0.85 mm isotropic voxel size) on a Philips 3T Achieva scanner, (Philips, Best, Netherlands) prior to the main experiment. The field-of-view was 245×245×208 mm, covering the whole brain.

#### Pre-measurements and source projections

Each experiment started with a set of pre-measurements required to determine the subject-specific stimulation criteria for phase-triggered TMS. First, the electrode positions of the EEG were marked on the individual MPRAGE image using neuronavigation. After that, the individual position within the left M1-HAND eliciting the highest MEP response for the right FDI muscle M1-HAND hot-spot was functionally determined [21] and marked on the individual MRI using neuronavigation. After that, the individual motor resting threshold and intensity to elicit MEPs of approximately 1 mV were determined using an in-house threshold-hunting algorithm based on parameter estimation by sequential testing, implemented in Python. The individual mu-rhythm was identified by recording EEG during alternating blocks of rest and continuous isometric abduction (right FDI muscle). Each condition lasted 30s and was repeated six times, during both conditions, the participants were instructed to keep their eyes open. Prior to further processing, the individual peak frequency of the mu-rhythm was determined by the peak of the mu-power spectral density (PSD) during the rest conditions within 7-13 Hz. The muband was the defined as the interval from 2Hz below the peak and 2 Hz above the peak. In cases where the lower frequency limit (peak-2Hz) was below 7 Hz the limit was set at 7 Hz resulting in a narrower frequency band in these cases. The individual PSD maps were used to determine the individual mu-rhythm power and the stimulation threshold of the phase-algorithm was set to stimulate only when the power within the individual mu-band exceeded the 75th percentile during the rest condition. In order to project the EEG data to source space we formed a lead field matrix informed by the participants structural MRI scan using the “dipoli” boundary element method implemented in Fieldtrip (version date: 2016-01-26), the source considered was a single dipole with radial orientation at the position of the functionally determined FDI muscle M1-HAND hot-spot in all participants.

#### Main experiment

Figure 2 illustrates the experimental design. Using the online readout of endogenous EEG activity, we targeted four phases (0°, 90°, 180°, 270°) in a frequency band covering a frequency range of 2Hz above and below individual peak mu frequency. Since the TMS pulse heavily disrupts EEG samples at the time immediately following stimulation, a trigger was set in 50% of all trials without delivering a TMS pulse. This enabled post-hoc evaluation of phase estimation performance in the non-stimulated trials, while phase-dependent effects of TMS on corticospinal excitability was evaluated in the 50% of trials with TMS. This setup resulted in a total of eight conditions (Stim0°, Stim90°, Stim180°, Stim270° and Trigger0°, Trigger90°, Trigger180°, Trigger270°). 60 trials were recorded per condition, resulting in a total of 480 trials. To ensure that sufficient time elapsed between two consecutive TMS pulses and to avoid any systematic interaction between TMS and mu-rhythm (e.g. phase-resetting, prolonged suppression of corticomotor excitability) the minimum inter-trial-interval (ITI) set by the algorithm was 2 s. Due to the mu-power and mu-stability criteria implemented in the algorithm and the 50% non-stimulation trials, the actual intervals between two TMS trials, the Inter-Stimulus Interval (ISI), was in practice much longer with a mean ISI across all individuals of 11.9 s. On a few occasions (∼ 0.5% of trials) the ISI exceeded 60 s in which case we did not consider the following MEP in the analysis.

**Figure 2:**
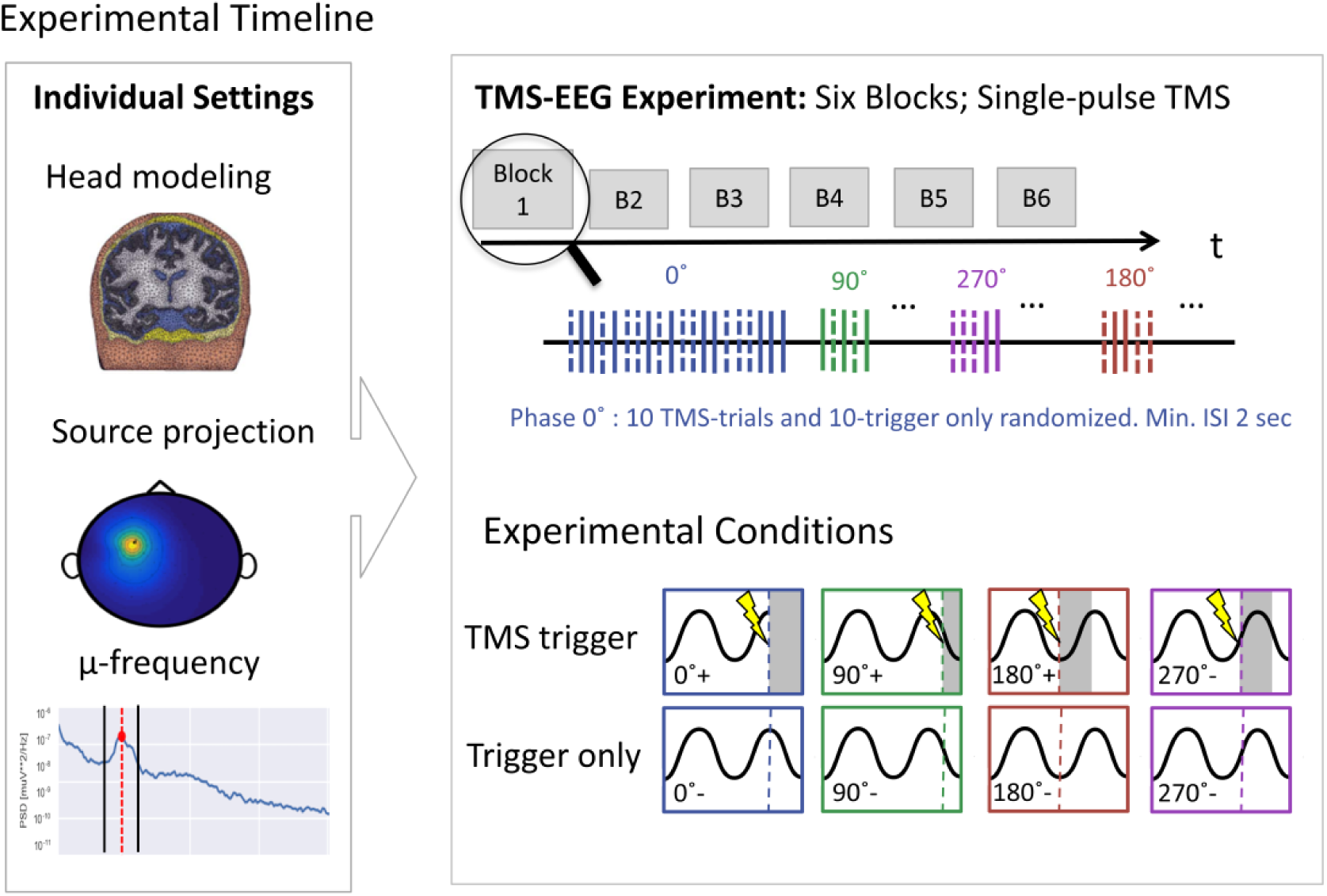
The experimental timeline: The first panel shows how the location and center frequency of the individual mu-rhythm were determined. The second panel illustrates the temporal structure of the main experiment and the eight conditions.

The experimental settings rendered the duration of the experiment quite long and hence we decided to split the main experiment in six shorter sessions to give participants short breaks of 1-5 minutes (80 trials per session, 10 trials per condition). In each session, the four phases were tested in blocks of 20 trials with 10 trials without or with TMS applied in a pseudo-random order. At the beginning of each block it was checked if the current stimulation intensity still elicited a MEP of 1 mV, if this was not the case the stimulator output was adjusted accordingly. Note that this never resulted in a change of more than 2% stimulator output. Average stimulator output was 73% ± 15% of the maximal stimulator output.

### Data Analysis

#### MEP analysis

The peak-to-peak amplitude of the MEP was determined trial-by-trial using an in-house developed python script. Trials that either displayed EMG activity > 50 uV in the 100 ms prior to stimulation onset or that were more than 2.5 standard derivations away from the individual MEP average were discarded from further analysis. On average 2.8 % of trials were excluded per individual participant due to these criteria.

#### Pre-stimulus power

Pre-stimulus power for each trial was estimated as the fraction of power within the mu-band based on a discrete Fourier transform of data in a window 500 ms prior to the stimulation.

#### Statistics

To test our main hypothesis, we investigated the population-averaged effects of phase triggered TMS on MEP amplitudes computing repeated measures ANOVA, with mu-phase (0°, 90°, 180°, 270°) as the independent within-subject variable, and the mean log-transformed MEP amplitude as dependent variable. Since analysis of mean MEP amplitude across subjects did not reveal any significant effect of mu-phase (see results), we decided to perform a more sensitive mixed-effects analysis incorporating mu power and the interval between two consecutive trials as additional factors in the statistical model. The linear mixed effects model included mu-phase (categorical – 0°,90°,180°,270°), mu-power (continuous) and the interval between two consecutive stimuli (ISI, continuous) as fixed effects, treating the participant factor as a random effect. Statistical analyses were performed using the lme4 package (Team RC 2018) within the R statistical software package (version 3.5.0; https://www.R-project.org). The significance threshold for null hypothesis testing was set at p<0.05.

We performed follow-up correlation analysis to test whether the individual phase-related differences in mean MEP amplitude were predicted by how strongly individual mu-activity was expressed in the left pericentral Rolandic cortex. To this end, we used the fraction of power within the mu-band detected during the pre-experiment resting EEG session as the independent variable, and the difference between the average MEP in two opposing phases (0° vs. 180° and 90° vs. 270° respectively).

## Results

### On-line phase-triggered EEG-TMS

On average, brain-state informed TMS targeted the intended phase (Figure 3). Analysis of the real-time triggered non-stimulated trials showed that the intended phase was targeted with a mean absolute error of 48.9° across all subjects and phases (Figure 3 b, for phase-specific errors). The average raw EEG-signal preceding the TMS trigger in each of the four conditions was also plotted. Targeting errors were symmetrically centered on the targeted phase. For individual phases the mean targeted phase and the mean absolute errors were as follows: Phase 0° = 5°±50°, Phase 90° = 78°±51°, Phase 180° = 177°± 52° and Phase 270° = 262°±48°. Despite the considerable inter-subject variability in the regional expression of pericentral alpha activity at rest, accuracy of the current algorithm is comparable with the performance of previously published solutions in pre-selected individuals with strong mu-power [34].

**Figure 3:**
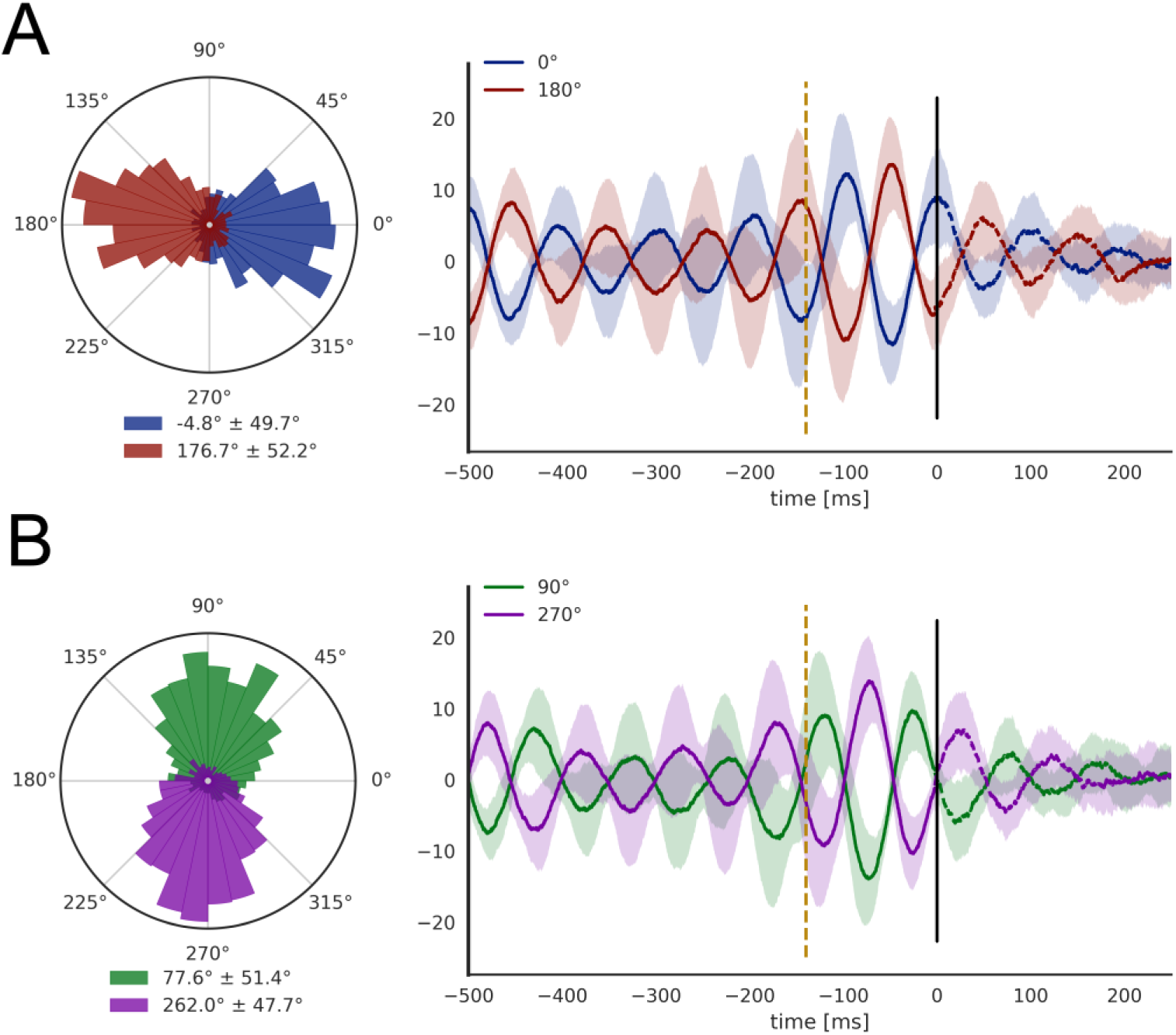
Performance of phase targeting. Panels A and B show phase targeting performance for 0° versus 180° and 90° versus 270° respectively. Circular phase histograms are presented on the left side of each panel. The histograms are calculated based on the non-stimulation trials in which phase is estimated using windows centered on the intended stimulation time. The graphs on the right side show the averaged pre-stimulus EEG activity prior to the estimated time of stimulation for stimulation and non-stimulation trials. The shaded areas cover the standard deviation across subjects. The part of the curve shortly before and after the estimated time point of stimulation issues dashed lines to indicate that this part of the curve only includes non-stimulation trials due to the large TMS artefact present for those trials.

### Motor Evoked Potentials

Mean MEP amplitude was 1.05±0.42 mV across all conditions and did not display a consistent phase-dependent variation (see figure 4). For individual phases the mean MEP amplitudes were as follows: Phase 0° = 1.05 mV, Phase 90° = 1.05 mV, Phase 180° = 1.01 mV and Phase 270° = 1.10 mV. Using the log-transformed mean MEP amplitudes as dependent variable, repeated-measures ANOVA revealed no significant effect of phase (F(3,39)=1.19; p=0.325). There were also no phase-related differences when directly comparing opposing phases (0 vs. 180 and 90 vs 270) using two-sided t-test (t_13_=1.50; p=0.157 and t_13_=-1.00; p=0.336). Figure 4A shows the mean individual as well as the mean group data for each target phase.

**Figure 4:**
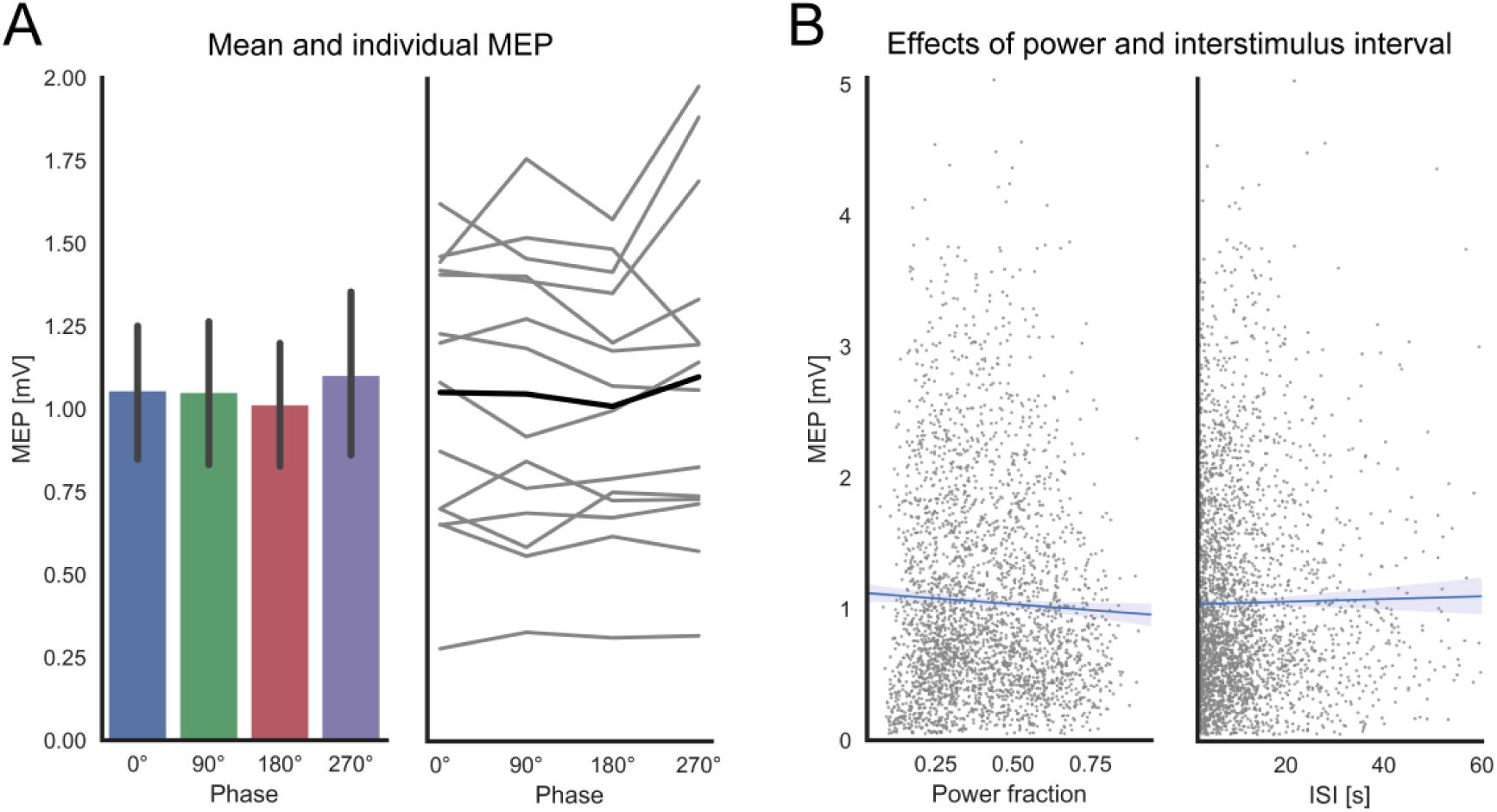
The effects of mu-phase, mu-power and inter-stimulus interval on the MEP amplitude. Panel A shows the grand averaged MEP data as a function of phase (lines indicating 95% confidence intervals) on the left and the individual MEP averages as a function of phase on the left. The black line indicates the mean. Panel B shows the correlation between individual MEP amplitudes (flexible model) and the pre-stimulus fraction of mu-power (left panel) and the ISIs (right panel). The blue lines are linear regression lines, with shaded areas indicating 95% bootstrapped confidence intervals.

A more comprehensive linear mixed-effects model treated mu-phase, mu-power, and ISI as fixed effects and participants as random effect also showed no significant main effect for mu-phase (χ^2^(3)= 3.21, p= 0.360), but main effects for mu-power (χ^2^(1)= 8.61, p< 0.003 and ISI (χ^2^(1)= 7.36, p =0.006). None of the interaction terms were significant (all p>0.2). The simple main effect of mu-power was due to a linear decrease in MEP amplitude with the level of mu-power at the time of TMS (Figure 4B). The simple main effect of ISI reflected a linear increase in MEP amplitude with longer intervals between consecutive TMS pulses (Figure 4B). Figure 4B shows that the effect size of the significant main effects (power and ISI) were very small in size given the large variability of individual MEPs.

We found no significant relationship between the magnitude of individual mu-rhythm expression during resting EEG epochs in the in preparatory EEG session and phase-related differences in mean MEP amplitude in the main experiment (Figure 5). The individual prominence of intrinsic mu-activity at rest did neither scale with the individual difference in MEP amplitude at mu-phase 0° vs. mu-phase 180° (t_13_=1.50, p=0.16) nor with the individual difference in MEP amplitude at mu-phase 90° vs. mu-phase 270° (t_13_=-1.00, p=0.34).

**Figure 5:**
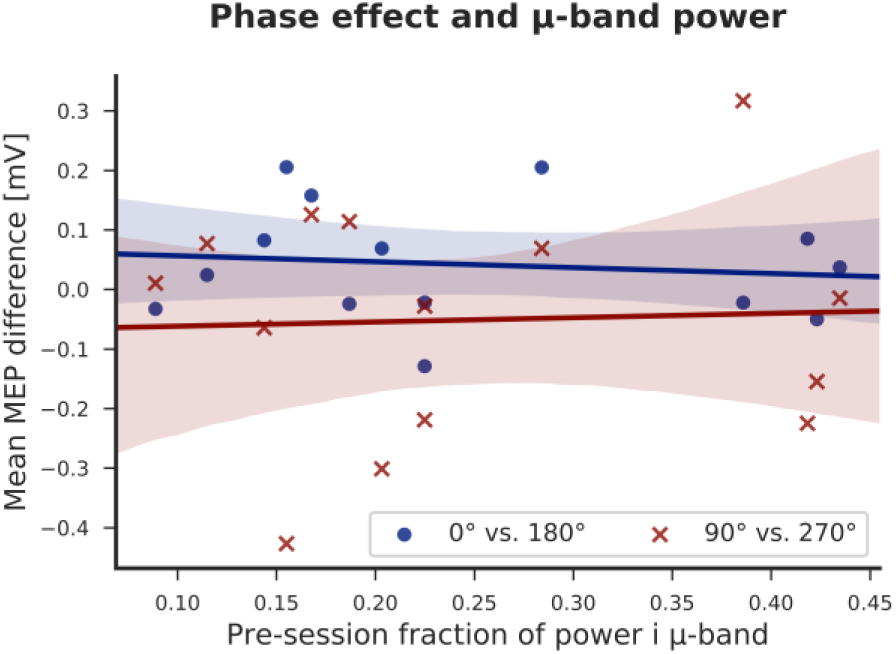
No significant relationship between prominence of mu-rhythm in the pre-session resting-EEG and phase-related differences in mean MEP amplitude can be seen for 0° vs. 180° (blue) nor for 90° vs 270° (red).

## Discussion

We demonstrate that phase-triggered TMS at four distinct phases of the ongoing mu-oscillations is technically feasible in non-preselected young volunteers. Contrary to our hypothesis, brain-state informed TMS targeting the peak and trough as well as the ascending and descending phase of ongoing mu-activity did not reveal any consistent modulatory effect of mu-phase on corticospinal excitability. Analysis of mean MEP amplitudes across subjects failed to reveal phase-related effects. A mixed effects analysis which also considers within subject variability also showed no effect of mu-phase on MEP amplitudes and yielded no interactions of mu-phase with mu-power or ISI. Instead, we were able to observe a weak but statistically significant effect of pre-stimulus mu-power and ISI, when including these factors as continuous variables in our mixed-model analysis. Mixed effects analysis revealed that higher mu-power resulted in slightly decreased MEP amplitudes while longer ISI intervals lead to slight increases in the MEP amplitude. Neither mu-power nor ISI interacted with oscillatory phase, and the size of these effects was small when considering the large variability of MEP amplitudes.

Our main finding that the phase of local ongoing mu-oscillations in the stimulated Rolandic cortex does not modulate MEP size is consistent with previous studies using post-hoc binning of single trials according to ongoing mu-phase at the time of stimulation. These studies found no or only very weak effects of mu-phase on MEP amplitude [24, 25, 27, 28, 31]. However, the most recent of these studies reported an interaction between pre-stimulus power and pre-stimulus phase when using a linear mixed-effects model [31]: TMS given at the trough of mu-oscillations elicited larger MEP amplitudes compared to TMS at the mu-peak when the pre-TMS power was high. The effect of phase was reversed during low-power trials during which MEP amplitudes evoked at the mu-peak were larger than MEP amplitudes evoked at mu-trough. No relationship between mu-phase and MEP amplitudes were present at medium levels of mu-power [31]. Our mixed-effect analysis confirmed the lack of a main effect of phase reported by Hussein et al., but did not reveal a consistent phase-power interaction in the mu-band. To explain this apparent discrepancy, it is important to point out that we only stimulated when individual mu-power was in the highest 2.5 percentiles and hence tested the effect of phase in brain states characterized by the maximal possible mu-expression in our participants. We did so, because we expected that a restriction of phase-triggered TMS to epochs with relatively strong mu-expression would yield the largest possible phase modulation, yet we did not detect a significant phase effect. Despite the relatively narrow power range, we also explored the possibility of a phase-power interaction in our data set, but did not find a significant phase-power interaction. Our negative finding is in agreement with intracranial recordings in humans, reporting no mu-phase-dependent modulations in neuronal firing in human somatosensory cortex [39].

Three recent studies, all performed by the same group also used a brain-state triggered EEG-TMS setup to investigate the influence of mu-phase on MEP amplitude [33-35]. In contrast with our negative findings, these studies showed significantly larger MEP amplitudes when TMS was given in the trough compared to the peak of the mu-oscillation. Several differences in the experimental approach may account for this discrepancy. In contrast to our study, participants were strongly pre-selected based on the magnitude of mu-power expressed in the resting-state EEG and more than 60% of all screened individuals were excluded because they did not meet the pre-set power criterion. The strict pre-selection of participants may impact the generalizability of their findings especially as the same group has demonstrated in subsequent work that the individual response to mu-phase triggered TMS is variable [33]. Using a bootstrapping approach to test the intra-individual phase-dependent modulation the authors showed that only one-third of the already preselected high-mu power participants demonstrated reliable intra-individual phase modulation. It may be possible that the reported modulation of MEP amplitude by mu-phase may be driven by a relatively limited subgroup of people. Hence, applying phase-dependent TMS to the general population, irrespective of baseline power, may further dilute the number of ‘responders’ and hence not result in significant effects at the group level. Of note, is that a post-hoc analysis of our data did not indicate that individuals with relative strong mu-power at baseline differed from those with relative weak mu-power. This observation argues against the notion that we might have found a phase relationship by applying a pre-selection based on baseline mu-power in the present study.

Additionally, the work by Schaworonkow et al. suggested that large numbers of trials (> 100 per condition) were needed to detect reliable phase modulation at the individual level. Due to the number of conditions and the relatively long ISI used in this study we did not have that many trial per conditions. While this may have limited our ability to detect very weak relationships between the MEP amplitude and mu-phase, we would argue that a modulatory effect only detectable by averaging more than 100 trials, might only marginally contribute to the inter-trial variability of the MEP.

Another potentially relevant difference between the present work and previous brain-state informed EEG-TMS studies is the inter-stimulus interval (ISI). In our study, the ISI between consecutive TMS stimuli lasted on average 11.9 seconds based on the power and stability criteria implemented in the phase-dependent triggering. The ISI was both significantly shorter and less variable in the work by Zrenner and coworkers, resulting in a quasi-repetitive stimulation at 0.5Hz [33, 34]. To explore if the temporal duration of the ISI effected the phase-modulation, we included the ISI as a variable in our statistical analysis. We were not able to detect any interactions between oscillatory phase and ISI, but found that shorter ISIs lead to lower MEP amplitudes, an effect that has been previously reported [40]. In fact, several studies have showed that continuous quasi-repetitive, or jittered application of supra-threshold TMS in the 0.5 - 0.3 Hz range can induce changes in the excitatory-inhibitory balance in M1 causing decrease in intracortical facilitation [41] as well as increased intracortical inhibition [42]. While some of these studies reported changes in the inhibitory/excitatory balance without changes of general MEP amplitude, newer studies suggest that also the MEP amplitude may be systematically modified by the inter-stimulus interval [43]. The long lasting effect of single TMS pulses can also be quantified by other physiological markers: Near Infrared Spectroscopy (NIRS) data can show a dip in oxy-hemoglobin following a TMS pulse that takes > 10 seconds to recover [44]. Taken together, this suggests that the stimulus interval has an effect on corticospinal excitability. We hypothesize that short ISIs above 0.2 HZ induced an “inhibitory brain state” in the targeted M1-HAND and its corticospinal output areas. This active state modulation introduced by a large number of repetitive suprathreshold TMS pulses might have triggered a mu-phase dependent [33, 34] and mu-power dependent modulation [36]. The mu-dependent effects that present in a TMS-induced inhibitory state might not be representative for the physiological mu-effects that can be detected in normal non-inhibited resting state by very low frequency TMS at highly variable ISIs – as used in the present study. Future studies are required to systematically assess the influence of ISI on mu-dependent fluctuations in corticospinal excitability.

While we found no effect of mu-phase, mixed-effect analysis showed that the level of mu-power expressed just before TMS had an effect on MEP amplitude. The negative correlation between high pre-stimulus mu-power and MEP amplitude in our data is consistent with previous work investigating interactions between oscillatory mu-power and corticospinal excitability, showing that higher levels of pre-stimulus mu-power are associated with lower MEP amplitudes [16, 22, 23]. Yet, the effect of mu-power was weak relative to overall variability of MEP amplitudes and showed a relatively flat regression slope. Further, the results cannot be generalized beyond the studied group of participants, as the detected effect was dependent on the inclusion of a random factor, modeling all data points of each participants. Interestingly, a recent state-informed EEG-TMS study, which used the mu-power as a criterion to trigger TMS, reported the reversed relationship. In that study, MEP amplitudes were larger, the higher pre-stimulus power [36]. A similar positive correlation between MEP amplitude and pre-stimulus mu-power is also evident in the supplementary material of a recent study in which TMS was triggered according to mu-phase [34]. It is likely that the relative short ISI in these studies may have impacted the general excitability level of M1-HAND and thereby introduced a paradoxical facilitation of corticospinal excitability by the intrinsically expressed mu power.

When considering all published data, it becomes obvious that the relationship between corticospinal excitability as probed with single-pulse MEP and the intrinsically expressed pericentral mu-rhythm is complex. Multiple intrinsic factors such as mu-phase or mu-power as well as extrinsic factors such as stimulation intensity, pulse configuration or ISI of TMS may result in complex interactions which might result in divergent findings. To complicate things further, cortical oscillations in the beta and gamma range have also been suggested to modulate corticospinal excitability [24, 29, 45-47]. In fact the influence of the sensorimotor beta rhythm has been suggested to be dominant within the Rolandic cortex [39, 48] and several studies have found associations between MEP amplitude and the pericentral beta rhythm [24, 29]. As the MEP amplitude is a compound measure of cortical and spinal excitability, also the fluctuations on coupling between cortical and spinal oscillations may be a potential modulator of MEP amplitudes and it has been suggested that cortico-spinal coupling in the beta range may have a greater influence on MEP amplitudes than intrinsic cortical oscillations [25, 28, 47].

From a methodological perspective, EEG-informed phase targeting poses an inherent challenge regarding the ability to optimally detect the relevant phase information. EEG is sensitive to electrical fields generated by radially or tangentially oriented dipoles. In this study, we have confined our source detection algorithm to radial dipoles as this allowed comparable phase weighting across participants which should give comparable sensitivity with the Hjorth montage used by Zrenner et al. (2018). Considering also tangential sources would complicate across-subject averaging, because differences in the algebraic sign of electrode weighting make averaging across participants difficult. However a tangential source might better represent the individual mu-source that displays mu-related fluctuations in MEP amplitude, because TMS induces a tangentially oriented electrical field in the cortex. These issues remain to be addressed in order to identify the best way of extracting the oscillatory EEG signal for phase-dependent TMS targeting of cortical oscillations.

*In conclusion,* phase-depended state-informed EEG-TMS is an interesting and promising tool for understanding the neurophysiological principles of cortical oscillations and their role in influencing neural excitability. The complex and yet unexplored underlying mechanisms still require a considerable amount of additional research before phase-dependent applications can reliably be used to decrease intra-individual variability in the response to TMS and provide a solid framework for boosting the effectiveness of current TMS-protocols.

## Acknowledgements

We would like to thank Chloe Chung, Syochi Tashiro and Borhan Javanmiri for help with the data collection. This work was supported by the Novo Nordisk Foundation Interdisciplinary Synergy Program 2014 [“Biophysically adjusted state informed cortex stimulation (BASICS); NNF14OC0011413]. HRS holds a clinical professorship in precision medicine at the Institute of Clinical Medicine, Faculty of Health and Medical Sciences, University of Copenhagen, sponsored by the Lundbeck Foundation.

## Competing Interests

Hartwig R. Siebner has received honoraria as speaker from Novartis, Denmark and Sanofi Genzyme, Denmark, as consultant from Sanofi Genzyme, Denmark, and as senior editor (NeuroImage) from Elsevier Publishers, Amsterdam, The Netherlands. He has received royalties as book editor from Springer Publishers, Stuttgart, Germany and has received research funding from Biogen-idec, Denmark. The other authors report no conflict of interests,

## References

[1] Maris E, Fries P, van Ede F. Diverse Phase Relations among Neuronal Rhythms and Their Potential Function. Trends Neurosci 2016;39(2):86–99.

[2] Bonnefond M, Kastner S, Jensen O. Communication between Brain Areas Based on Nested Oscillations. eNeuro 2017;4(2).

[3] Klimesch W. The frequency architecture of brain and brain body oscillations: an analysis. Eur J Neurosci 2018;48(7):2431–53.

[4] Jensen O, Mazaheri A. Shaping functional architecture by oscillatory alpha activity: gating by inhibition. Frontiers in human neuroscience 2010;4:186.

[5] Zumer JM, Scheeringa R, Schoffelen JM, Norris DG, Jensen O. Occipital alpha activity during stimulus processing gates the information flow to object-selective cortex. PLoS Biol 2014;12(10):e1001965.

[6] Herring JD, Thut G, Jensen O, Bergmann TO. Attention Modulates TMS-Locked Alpha Oscillations in the Visual Cortex. J Neurosci 2015;35(43):14435–47.

[7] de Pesters A, Coon WG, Brunner P, Gunduz A, Ritaccio AL, Brunet NM, et al. Alpha power indexes task-related networks on large and small scales: A multimodal ECoG study in humans and a non-human primate. Neuroimage 2016;134:122–31.

[8] Jensen O, Gips B, Bergmann TO, Bonnefond M. Temporal coding organized by coupled alpha and gamma oscillations prioritize visual processing. Trends Neurosci 2014;37(7):357–69.

[9] Gips B, van der Eerden JP, Jensen O. A biologically plausible mechanism for neuronal coding organized by the phase of alpha oscillations. Eur J Neurosci 2016;44(4):2147–61.

[10] Romei V, Rihs T, Brodbeck V, Thut G. Resting electroencephalogram alpha-power over posterior sites indexes baseline visual cortex excitability. Neuroreport 2008;19(2):203–8.

[11] Romei V, Brodbeck V, Michel C, Amedi A, Pascual-Leone A, Thut G. Spontaneous fluctuations in posterior alpha-band EEG activity reflect variability in excitability of human visual areas. Cereb Cortex 2008;18(9):2010–8.

[12] Mathewson KE, Lleras A, Beck DM, Fabiani M, Ro T, Gratton G. Pulsed out of awareness: EEG alpha oscillations represent a pulsed-inhibition of ongoing cortical processing. Frontiers in psychology 2011;2:99.

[13] Bruers S, VanRullen R. Alpha Power Modulates Perception Independently of Endogenous Factors. Frontiers in neuroscience 2018;12:279.

[14] Pfurtscheller G, Neuper C, Andrew C, Edlinger G. Foot and hand area mu rhythms. Int J Psychophysiol 1997;26(1-3):121–35.

[15] Linkenkaer-Hansen K, Nikulin VV, Palva S, Ilmoniemi RJ, Palva JM. Prestimulus oscillations enhance psychophysical performance in humans. J Neurosci 2004;24(45):10186–90.

[16] Anderson KL, Ding M. Attentional modulation of the somatosensory mu rhythm. Neuroscience 2011;180:165–80.

[17] Ai L, Ro T. The phase of prestimulus alpha oscillations affects tactile perception. Journal of Neurophysiology 2014;111:1300–7.

[18] Forschack N, Nierhaus T, Muller MM, Villringer A. Alpha-Band Brain Oscillations Shape the Processing of Perceptible as well as Imperceptible Somatosensory Stimuli during Selective Attention. J Neurosci 2017;37(29):6983–94.

[19] Haegens S, Nacher V, Luna R, Romo R, Jensen O. alpha-Oscillations in the monkey sensorimotor network influence discrimination performance by rhythmical inhibition of neuronal spiking. Proc Natl Acad Sci U S A 2011;108(48):19377–82.

[20] Hallett M. Transcranial magnetic stimulation: a primer. Neuron 2007;55(2):187–99.

[21] Groppa S, Oliviero A, Eisen A, Quartarone A, Cohen LG, Mall V, et al. A practical guide to diagnostic transcranial magnetic stimulation: report of an IFCN committee. Clin Neurophysiol 2012;123(5):858–82.

[22] Zarkowski P, Shin CJ, Dang T, Russo J, Avery D. EEG and the variance of motor evoked potential amplitude. Clinical EEG and neuroscience 2006;37(3):247–51.

[23] Sauseng P, Klimesch W, Gerloff C, Hummel FC. Spontaneous locally restricted EEG alpha activity determines cortical excitability in the motor cortex. Neuropsychologia 2009;47(1):284–8.

[24] Maki H, Ilmoniemi RJ. EEG oscillations and magnetically evoked motor potentials reflect motor system excitability in overlapping neuronal populations. Clin Neurophysiol 2010;121(4):492–501.

[25] van Elswijk G, Maij F, Schoffelen JM, Overeem S, Stegeman DF, Fries P. Corticospinal beta-band synchronization entails rhythmic gain modulation. J Neurosci 2010;30(12):4481–8.

[26] Takemi M, Masakado Y, Liu M, Ushiba J. Event-related desynchronization reflects downregulation of intracortical inhibition in human primary motor cortex. J Neurophysiol 2013;110(5):1158–66.

[27] Berger B, Minarik T, Liuzzi G, Hummel FC, Sauseng P. EEG oscillatory phase-dependent markers of corticospinal excitability in the resting brain. BioMed research international 2014;2014:936096.

[28] Keil J, Timm J, Sanmiguel I, Schulz H, Obleser J, Schonwiesner M. Cortical brain states and corticospinal synchronization influence TMS-evoked motor potentials. J Neurophysiol 2014;111(3):513–9.

[29] Schulz H, Ubelacker T, Keil J, Muller N, Weisz N. Now I am ready-now i am not: The influence of pre-TMS oscillations and corticomuscular coherence on motor-evoked potentials. Cereb Cortex 2014;24(7):1708–19.

[30] Iscan Z, Nazarova M, Fedele T, Blagovechtchenski E, Nikulin VV. Pre-stimulus Alpha Oscillations and Inter-subject Variability of Motor Evoked Potentials in Single- and Paired-Pulse TMS Paradigms. Frontiers in human neuroscience 2016;10:504.

[31] Hussain SJ, Claudino L, Bonstrup M, Norato G, Cruciani G, Thompson R, et al. Sensorimotor Oscillatory Phase-Power Interaction Gates Resting Human Corticospinal Output. Cereb Cortex 2018.

[32] Bergmann TO, Molle M, Schmidt MA, Lindner C, Marshall L, Born J, et al. EEG-guided transcranial magnetic stimulation reveals rapid shifts in motor cortical excitability during the human sleep slow oscillation. J Neurosci 2012;32(1):243–53.

[33] Schaworonkow N, Triesch J, Ziemann U, Zrenner C. EEG-triggered TMS reveals stronger brain state-dependent modulation of motor evoked potentials at weaker stimulation intensities. Brain Stimul 2018.

[34] Zrenner C, Desideri D, Belardinelli P, Ziemann U. Real-time EEG-defined excitability states determine efficacy of TMS-induced plasticity in human motor cortex. Brain Stimul 2018;11(2):374–89.

[35] Stefanou MI, Desideri D, Belardinelli P, Zrenner C, Ziemann U. Phase Synchronicity of mu-Rhythm Determines Efficacy of Interhemispheric Communication Between Human Motor Cortices. J Neurosci 2018;38(49):10525–34.

[36] Thies M, Zrenner C, Ziemann U, Bergmann TO. Sensorimotor mu-alpha power is positively related to corticospinal excitability. Brain Stimul 2018.

[37] Safeldt M, Tomasevic L, Karabanov A, Siebner H K M. Towards brain-state dependent transcranial magnetic stimulation: Targeting the phase of oscillatory neocortical activity with singe-pulse TMS. Brainstimulation 2017.

[38] Frigo M, Johnson SG. The Design and Implementation of FFTW3. Proceedings of the IEEE 2005;93(2):216–31.

[39] Miller KJ, Hermes D, Honey CJ, Hebb AO, Ramsey NF, Knight RT, et al. Human motor cortical activity is selectively phase-entrained on underlying rhythms. PLoS Comput Biol 2012;8(9):e1002655.

[40] Vaseghi B, Zoghi M, Jaberzadeh S. Inter-pulse Interval Affects the Size of Single-pulse TMS-induced Motor Evoked Potentials: A Reliability Study. Basic Clin Neurosci 2015;6(1):44–51.

[41] Gorsler A, Baumer T, Weiller C, Munchau A, Liepert J. Interhemispheric effects of high and low frequency rTMS in healthy humans. Clin Neurophysiol 2003;114(10):1800–7.

[42] Cincotta M, Borgheresi A, Gambetti C, Balestrieri F, Rossi L, Zaccara G, et al. Suprathreshold 0.3 Hz repetitive TMS prolongs the cortical silent period: potential implications for therapeutic trials in epilepsy. Clin Neurophysiol 2003;114(10):1827–33.

[43] Julkunen P, Saisanen L, Hukkanen T, Danner N, Kononen M. Does second-scale intertrial interval affect motor evoked potentials induced by single-pulse transcranial magnetic stimulation? Brain Stimul 2012;5(4):526–32.

[44] Thomson RH, Maller JJ, Daskalakis ZJ, Fitzgerald PB. Blood oxygenation changes resulting from trains of low frequency transcranial magnetic stimulation. Cortex 2012;48(4):487–91.

[45] Lepage JF, Morin-Moncet O, Beaule V, de Beaumont L, Champoux F, Theoret H. Occlusion of LTP-like plasticity in human primary motor cortex by action observation. PLoS One 2012;7(6):e38754.

[46] Kraus D, Naros G, Bauer R, Khademi F, Leao MT, Ziemann U, et al. Brain State-Dependent Transcranial Magnetic Closed-Loop Stimulation Controlled by Sensorimotor Desynchronization Induces Robust Increase of Corticospinal Excitability. Brainstimulation 2016;accepted

[47] Khademi F, Royter V, Gharabaghi A. Distinct Beta-band Oscillatory Circuits Underlie Corticospinal Gain Modulation. Cereb Cortex 2018;28(4):1502–15.

[48] Groppe DM, Bickel S, Keller CJ, Jain SK, Hwang ST, Harden C, et al. Dominant frequencies of resting human brain activity as measured by the electrocorticogram. Neuroimage 2013;79:223–33.

